# Impact of regulatory variation across human iPSCs and differentiated cells

**DOI:** 10.1101/091660

**Authors:** Nicholas E. Banovich, Yang I. Li, Anil Raj, Michelle C. Ward, Peyton Greenside, Diego Calderon, Po Yuan Tung, Jonathan E. Burnett, Marsha Myrthil, Samantha M. Thomas, Courtney K. Burrows, Irene Gallego Romero, Bryan J. Pavlovic, Anshul Kundaje, Jonathan K. Pritchard, Yoav Gilad

## Abstract

Induced pluripotent stem cells (iPSCs) are an essential tool for studying cellular differentiation and cell types that are otherwise difficult to access. We investigated the use of iPSCs and iPSC-derived cells to study the impact of genetic variation across different cell types and as models for studies of complex disease. We established a panel of iPSCs from 58 well-studied Yoruba lymphoblastoid cell lines (LCLs); 14 of these lines were further differentiated into cardiomyocytes. We characterized regulatory variation across individuals and cell types by measuring gene expression, chromatin accessibility and DNA methylation. Regulatory variation between individuals is lower in iPSCs than in the differentiated cell types, consistent with the intuition that developmental processes are generally canalized. While most cell type-specific regulatory quantitative trait loci (QTLs) lie in chromatin that is open only in the affected cell types, we found that 20% of cell type-specific QTLs are in shared open chromatin. Finally, we developed a deep neural network to predict open chromatin regions from DNA sequence alone and were able to use the sequences of segregating haplotypes to predict the effects of common SNPs on cell type-specific chromatin accessibility.

## Main Text

The differentiation of iPSCs into a wide variety of cell types has become an essential tool for many research applications, and holds great promise in therapeutics [1]. One potentially important application is as a tool for studying the effects of common variation in cell types that are otherwise difficult to obtain.

In recent years it has become clear that much of the genetic basis of complex traits is due to common variants that affect cell type-specific gene regulation in relevant tissues and cell types [2, 3]. Consequently, numerous groups have studied variation in gene expression across panels of individuals in cell lines, in easily accessible primary tissue samples such as blood or skin [4, 5] or in post-mortem tissues [6]. However many important cell types cannot be obtained from adult post mortem samples; moreover, such tissues are unsuited for functional studies and perturbations that require living cells.

To explore the utility of an iPSC-based alternative model system, we generated a panel of iPSCs from 58 well-characterized Yoruba LCLs. Briefly, LCLs were reprogrammed using a previously-described episomal approach [7]. After a week in suspension, culture cells were seeded onto a layer of gelatin and mouse embryonic fibroblasts. A single colony was obtained from each line and passaged for ten weeks before final characterization, conversion to feeder-free growth, and collection. Pluripotency and stability were confirmed for each line (Supplementary Materials). This panel represents the largest stock of characterized non-European iPSCs to date and is available to other researchers, complementing parallel efforts in Europeans [8] (see Data Accession in Supplementary Materials).

To study gene regulation in iPSCs, we assayed three molecular phenotypes: mRNA expression (using RNA-seq; n=58), chromatin accessibility (ATAC-seq; n=57), and DNA methylation levels (EPIC arrays; n=58). We differentiated 14 iPSC lines into iPSC-derived cardiomyocytes (iPSC-CMs; Supplementary Materials) and collected RNA-seq and ATAC- seq from the 14 iPSC-CMs (Figure 1A). We analyzed these newly collected data jointly with data previously collected from the same Yoruba LCLs (we also added new ATAC-seq data for 20 of the LCLs).

**Figure 1:**
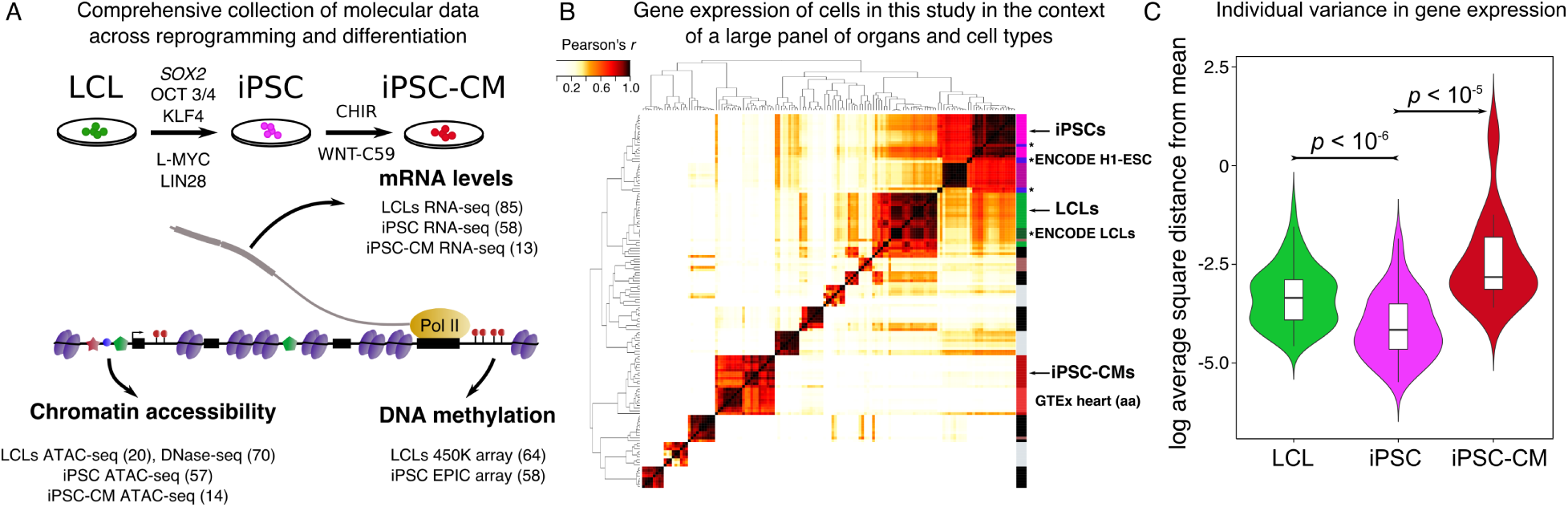
Systematic measurements of molecular phenotypes across reprogramming and differentiation. **(A)** Summary of data collection. **(B)** Correlation matrix of gene expression from our samples and samples from ENCODE (*) and GTEx. Our LCL samples cluster most closely with LCLs samples from ENCODE, while our iPSCs and iPSC-CM lines cluster most closely with H1- ESC (ENCODE) and heart (GTEx) respectively. Dark purple: GTEx bone marrow. **(C)** Violin plots representing per individual log_2_ of the average square distance from the mean (Supplementary Materials) for iPSC, LCL, and iPSC-CM gene expression levels. Plots for chromatin accessibility, and DNA methylation levels are shown in Supplementary Figure 1.

In the context of RNA-seq data from a larger panel of tissues and cell types from GTEx and ENCODE, respectively, gene expression data from our LCLs cluster most closely with data from ENCODE LCLs, as expected. Similarly, gene expression data from our iPSCs cluster with data from H1 embryonic stem cell lines from ENCODE, and data from our iPSC-CMs cluster most closely with gene expression data from GTEx heart tissues (atrial appendages) (Figure 1B, Supplementary Materials). Thus, our cultured cells broadly recapitulate expected regulatory patterns.

Notably, we observed that chromatin accessibility, gene expression, and DNA methylation levels were all more homogenous between individuals in iPSCs than in LCLs or iPSC-CMs (*p* < 10^−5^, Figure 1C, Supplementary Figure 1). This is consistent with the notion that developmental processes are canalized [9] and that regulatory states in embryonic cells are tightly controlled.

After examining overall properties in our data, we sought to characterize the effect of genetic variation on gene regulation. While there have been numerous multi-tissue studies of expression, our data provide the first opportunity to study QTLs for open chromatin in a cell type other than LCLs.

We first analyzed data from each cell type independently. We identified thousands of putatively cis genetic associations with all three regulatory phenotypes at 10% FDR (Supplementary Materials; Table S3). Despite the observation that regulatory phenotypes are associated with lower inter-individual variation in iPSCs compared to LCLs, we found similar or greater numbers of expression QTLs (eQTLs) in iPSCs when sample sizes are matched across cell types (e.g. 1,441 eQTLs in iPSCs versus 1,168 in LCLs using 58 individuals). In addition, using WASP [11], we identified 517 eQTLs and 3,989 chromatin accessibility QTLs (caQTLs) in the small sample of differentiated iPSC-CMs. In general, we observed a high degree of QTL sharing between cell types. We found 71% to 91% overlap in eQTLs between iPSCs and LCLs, using an estimate of sharing that accounts for incomplete power of the replication tests (Storey’s π_0_) (Supplementary Figure 3). The proportion of sharing is lower when considering iPSC-CMs (Supplementary Figure 3), as expected given the difference in sample size.

The high sharing of regulatory QTLs across cell types notwithstanding, we asked about the mechanisms by which some genetic variants affect gene regulation in one cell type with no detectable effect in other cell types. This is of interest given that disease-associated variants are enriched in open chromatin and particularly in cell type-specific open chromatin [13]. Indeed, as might be expected, we found that the iPSC-specific caQTLs we identified (Supplementary Materials) had larger effects on gene expression levels in iPSCs than did LCL-specific caQTLs, and conversely (*p* = 0.01, *p* = 4.7 × 10^−5^, respectively, Fisher’s exact test, Figure 2A). Some of these cell-type-specific caQTLs are located quite far from the gene they regulate (e.g. 50kb or more), and likely function by affecting distal enhancer or promoter elements (Supplementary Figure 4).

**Figure 2:**
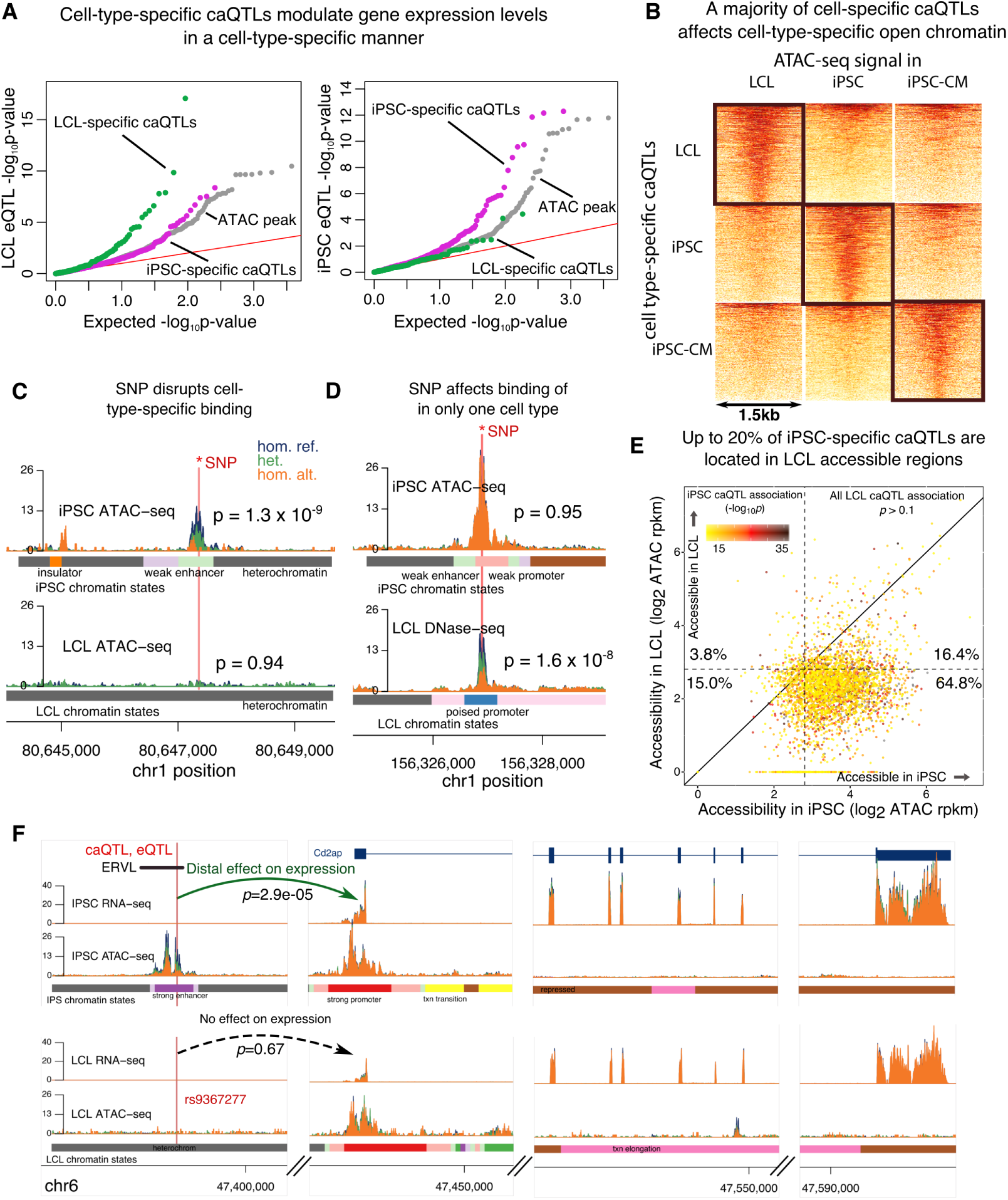
Mechanisms of cell-type-specific regulatory variation. **(A)** QQ-plot of LCL and iPSC eQTL signal conditioned on LCL- and iPSC-specific caQTLs. Higher enrichment of LCL (iPSC) eQTLs among LCL (iPSC) caQTLs links cell-type-specific regulation of chromatin accessibility to cell-type-specific regulation of gene expression. **(B)** Chromatin accessibility signal around cell-specific caQTLs in corresponding cell types (black rectangles), and in other cell-types. A lack of accessibility in other cell-types suggests that cell-specific caQTLs often affect cell-specific accessible regions, e.g. example (C). **(C-D)** Examples of cell-type-specific regulatory effects of genetic variation. SNP is correlated with accessibility of an iPSC-specific open chromatin region in iPSCs only (C), or of a non-specific open chromatin region in LCLs only (D). **(E)** Scatter plot of iPSC and LCL chromatin accessibility at iPSC-specific caQTLs. About 20% of iPSC-specific caQTLs are accessible in LCLs. Plot of LCL-specific caQTLs in Supplementary Figure 6. **(F)** Example of an iPSC-specific caQTL that is also an iPSC-specific eQTL. SNP rs9367277 is associated with both chromatin accessibility of a strong enhancer and with expression of the *Cd2ap* gene in iPSCs. Interestingly, rs9367277 lies in a transposable element of the ERVL family, which are preferentially activated in embryonic stem cells [10].

We further asked about the mechanisms by which genetic variants affect chromatin accessibility broadly, in multiple cell-types, or specifically in a single cell-type. As expected, caQTLs that are shared across cell-types lie within regulatory regions that are accessible in all cell-types, and likely affect the DNA binding of the same factors (Supplementary Figure 6). In contrast, most cell type-specific caQTLs lie in regions that are accessible in the affected cell type, but show little or no accessibility in the other cell types (Figure 2B,C). Thus, most (>70%) cell type-specific caQTLs can be explained simply by cell type-specific regulatory activity (Figure 2B).

While the notion that cell type-specific caQTLs can often be explained by cell type-specific chromatin activity is quite intuitive, we also found numerous regions that were accessible in multiple cell types, but with a regulatory effect in a single cell-type only (Figure 2D,F; Table S6 for a list). In fact, up to 20% of cell-type-specific caQTLs are accessible in multiple cell-types (Figure 2E). This observation is consistent with the idea that multiple DNA-binding factors may affect chromatin activity at the same locus by binding to distinct but nearby motifs [12].

Our observations that cell type-specific open chromatin regions can often explain contrasting effects of genetic variants in different cell types motivated us to explore the sequence features underlying differences in chromatin activity across cell types. In particular, we aimed to identify DNA sequences that could predict cell type-specific effects of regulatory variants. We investigated the use of machine learning models to predict the chromatin activity of regulatory elements across our three cell types using DNA sequence only [13, 14, 15, 16]. We developed a four-layered neural network architecture, OrbWeaver, to predict cell type-specific chromatin accessibility of 500bp windows centered at a regulatory locus (Figure 3A, Supplementary Figure 7). In contrast to popular approaches that learn all the parameters of the neural network *de novo*, we used log-transformed position weight matrices (PWMs) of 1, 320 human transcription factors (Supplementary Materials) [17, 18] as the first layer of OrbWeaver. As training input, we used 282,088 loci that were identified as accessible in at least one of the three cell types. When testing our predictions on a held-out dataset of 7,151 loci, we achieved high accuracies in all three cell types: iPSC (AUC = 0.96), LCL (AUC = 0.90), and iPSC-CM (AUC = 0.91) (Fig. 3B; see Supplementary Figure 8 for precision recall results). We found that the use of transcription factor PWMs as the first layer of OrbWeaver yielded higher predictive accuracies with a simpler neural network architecture than with a more complex architecture that did not use transcription factor PWMs (Supplementary Figure 8).

**Figure 3:**
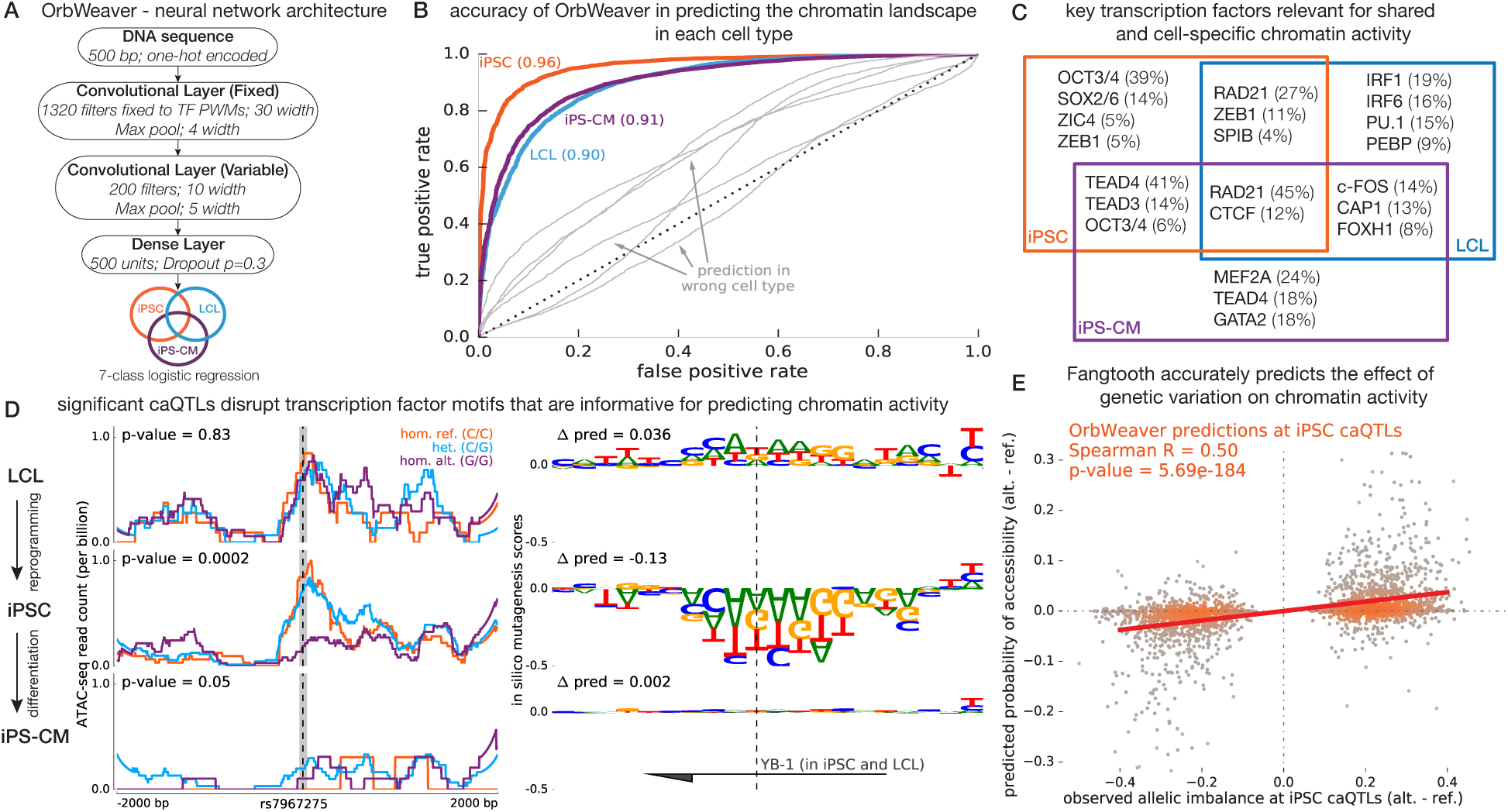
Predicting chromatin activity from sequence using deep neural networks. **(A)** OrbWeaver is a 4-layered neural network where the parameters of the first convolutional layer are fixed to known position weight matrices of human transcription factors. The activation function used in each of the convolutional and dense layers is the Rectified Linear Unit (ReLU). **(B)** The OrbWeaver model for one cell type poorly predicts open chromatin in other cell types (gray), highlighting that the model captures cell type-specific regulatory elements. **(C)** Transcription factors important for each locus were identified using DeepLIFT scores; this panel illustrates the top key TFs for each of the 7 categories of chromatin activity and the fraction of loci explained by them. **(D)** An example of a locus that is open in iPSCs and LCLs but was identified to be an iPSC-specific caQTL. The subpanels on the left show the raw ATAC-seq signal in each cell type stratified by genotype of the most significant SNP of the iPSC caQTL. The subpanels on the right show the marginal change in OrbWeaver predictions due to mutating the reference base at each position to an alternate base. The sequence shown corresponds to the shaded portion on the left subpanels and the reported Δpred values correspond to the change between alleles of the most significant SNP. The TF important for this locus as identified by DeepLIFT is YB-1, a factor highly expressed in all three cell types. **(E)** Scatter plot comparing the observed allelic imbalance at iPSC caQTLs, estimated by WASP, and the predicted difference in median chromatin activity between haplotypes tagged by the two alleles of the causal SNP. Note that the OrbWeaver model was learned using the reference genome sequence alone and had no information regarding genetic variation in the population when learning the model parameters.

To identify transcription factors that help predict the shared and cell type-specific regulatory activity across loci, we computed DeepLIFT scores [19] with respect to each filter in the first convolutional layer. Among 1,320 factors for which we had PWMs, the factor with the highest score for a given locus was assigned to be the most important factor for explaining the chromatin activity of said locus. Aggregating the key factor across all loci, we recovered transcription factors that are known to drive cell type-specific chromatin activity (Fig. 3C), and identified several additional factors that are putatively important for cell type-specific gene regulation (Table S6). Notably, nearly 40% of iPSC-specific open chromatin loci could be explained by the OCT3/4 motif alone. In LCLs and iPSC-CM, a larger number of TFs are needed to explain the same fraction of cell type-specific open chromatin loci. This observation is consistent with the higher predictive accuracy achieved for iPSCs compared to LCLs and iPSC-CMs, even with simpler neural network models (Supplementary Figure 8), and suggests that fewer trans-acting factors establish the chromatin landscape in embryonic cells than in somatic cells.

Given our ability to predict cell type-specific chromatin activity genome-wide from DNA sequence alone, we reasoned that OrbWeaver could also predict cell type-specific effects of SNPs on chromatin activity (Figure 3D). Prediction of SNP effects on gene regulation, especially in specific cell types, is a challenging problem, but is an essential task for future interpretation of personal genomes. Starting with iPSC caQTLs, we found that OrbWeaver predictions track the observed allelic imbalance ratio with a correlation of 0.50 (*p* = 6x 10^−184^; Figure 3E). Considering all tested SNPs in open chromatin peaks (the majority of which presumably have no true effect on chromatin accessibility) the correlation is more modest, though highly significant (iPSC correlation 0.12; *p* < 10^−308^). Notably, our ability to predict caQTL effects in one cell type is drastically reduced when using our model for another cell-type (Supplementary Figure 9), indicating that our model has high cell type-specificity. Altogether these findings demonstrate our ability to identify trans-acting elements driving cellular differences in chromatin accessibility and, more importantly, to predict effects of genetic variation in a cell type-specific manner.

Ultimately, the iPSCs and their differentiated cells may be valuable for developing a variety of models of human disease, provided that cultured differentiated cells are an effective system with which to model gene regulation in the corresponding primary tissue. We evaluated the fidelity of iPSC-CMs as a model for heart tissues and heart-related diseases. As discussed above, gene expression from iPSC-CMs most closely resembles those in GTEx heart samples. Furthermore, eQTLs detected in our iPSC-CMs are most enriched with eQTLs identified in GTEx heart tissues (left ventricle) (Supplementary Figure 2). We used a polygenic method (Supplementary Materials) to identify enrichments of GWAS signals associated with genes whose expression shows cell type specificity. Genes more specifically expressed in iPSC-CMs are enriched for signals from GWAS for body mass index (BMI), coronary artery disease (CAD), and myocardial infarction (MI), while genes more specifically expressed in LCLs are enriched for signals from GWAS for multiple sclerosis (MS), and rheumatoid arthritis (RA) (Figure 4A).

**Figure 4:**
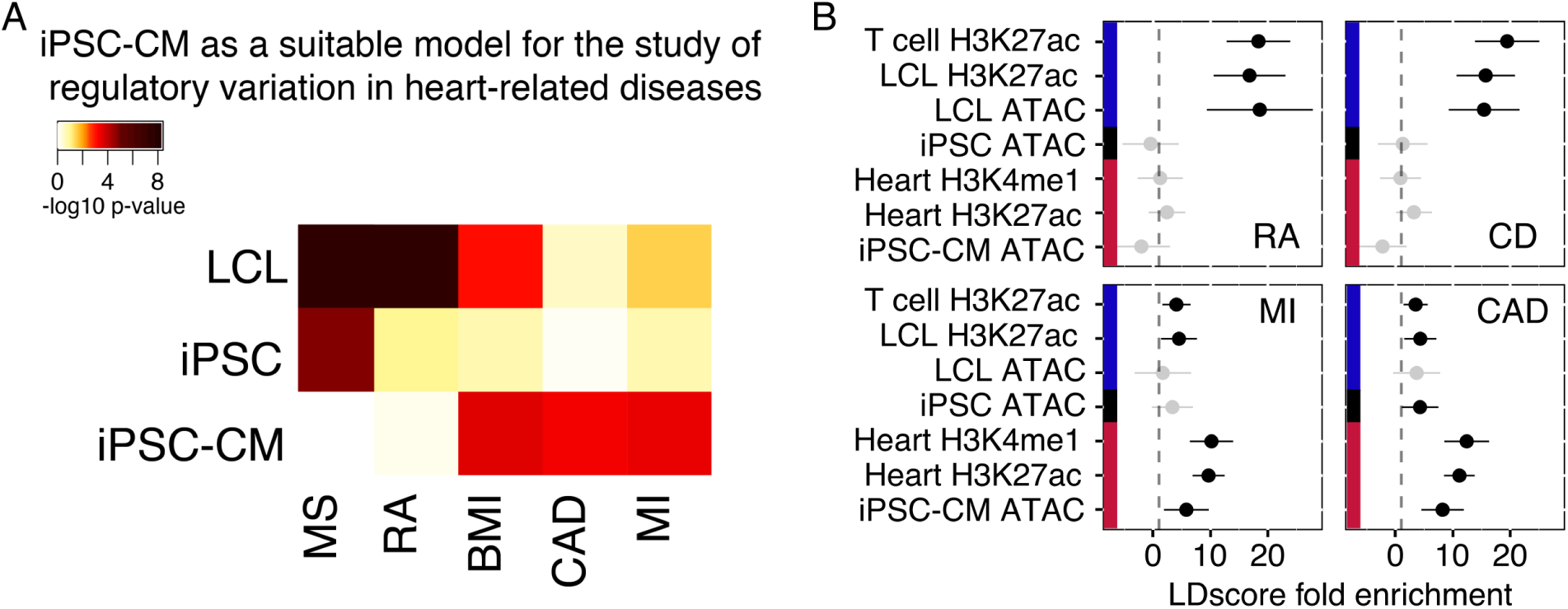
Modeling complex disease using iPSC-derived cells. **(A)** Heatmap of enrichment p- values of GWAS signals near genes with cell type-specific expression (Supplementary Materials). **(B)** Enrichments of SNPs associated with four different diseases in different partitions of the genome (computed using LDscore regression; point estimates ±95% confidence intervals). In both analyses, the autoimmune traits (multiple sclerosis (MS) or Crohn’s disease (CD) and rheumatoid arthritis (RA)) show enrichment near genes and chromatin that are more active in LCLs, and the heart-related traits (coronary artery disease (CAD) and myocardial infarction (MI)) are enriched in iPSC-CM active regions.

We also used stratified LD score regression [3] to estimate enrichment of heritability explained by GWAS signal within open chromatin in the different cell types (Figure 4B). We found that heritability explained by SNPs in LCL and iPSC-CM ATAC-seq peaks explained more heritability in autoimmune and heart-related diseases, respectively (all enrichment *p* ≤ 10^−2^). These observations suggest that cellular reprogramming followed by differentiation is a promising strategy to generate models of complex disease for which primary tissues are difficult to obtain.

In summary, we have established a unique resource of 58 fully characterized iPSC lines. These lines reprogrammed from LCLs obtained from Yoruba individuals originally collected as part of the HapMap project, represent the largest panel of iPSCs from individuals of African ancestry. We believe this resource will be of great value. In particular, future studies using this panel of iPSCs will be able to assay dynamic gene regulation by characterizing gene expression during differentiation, in multiple cell types from the same individuals, and in terminally differentiated cell types subjected to experimental perturbations. The move toward dynamic studies of gene regulation in disease relevant tissues will help to elucidate mechanisms underlying complex disease that were previously difficult or impossible to study. The research presented here is a first step towards this goal.

## Acknowledgments

We thank members of the Pritchard and Gilad Labs for helpful discussions. This work was supported by NIH grants GM007197, AG 044948,MH084703, MH101825, HG007036, CA149145, and HL092206; by a Center for Computational, Evolutionary and Human Genomics Fellowship; by a EMBO Long-Term Fellowship (ALTF 751-2014) and Marie Curie Actions; and by the Howard Hughes Medical Institute. All data have been deposited in the Gene Expression Omnibus (**www.ncbi.nlm.nih.gov/geo/**) under accession no. GSE89895; other accession numbers can be found in Table S7.

